# Stomatal maturomics: identifying genes regulating guard cell maturation and function formation from single-cell transcriptomes

**DOI:** 10.1101/2023.11.05.565663

**Authors:** Yuming Peng, Yi Liu, Yifan Wang, Yue Qin, Shisong Ma

## Abstract

Stomata play critical roles in gas exchange and immunity to pathogens. While many genes regulating early stomatal development up to the production of young guard cells (GCs) have been described in Arabidopsis, much less is known about how young GCs develop into mature functional stomata. Here we performed a maturomics study on stomata, with “maturomics” defined as omics analysis of the maturation process of a tissue or organ. We developed an integrative scheme to analyze three public stomata-related single-cell RNA-seq datasets and identified a list of 586 genes that were specifically up-regulated in all three datasets during stomata maturation and function formation. The list, termed sc_586, is enriched with known regulators of stomatal maturation and functions. We selected two candidate G2-like TFs genes, *MYS1* and *MYS2*, from the list to investigate their roles in stomata. Our results showed that these two genes redundantly regulate the size and hoop rigidity of mature GCs, and their double mutations caused mature GCs to have severe defects in regulating their stomatal apertures. Our analysis thus provides a valuable gene list for studying GC maturation and function formation.

## Introduction

Stomata are epidermal pores formed by pairs of guard cells on plant leaf surfaces, which regulate gas and water exchange between plants and the outside environment as well as immunity to pathogens. In Arabidopsis, stomatal development starts from a subset of protodermal cells termed meristemoid mother cells (MMCs) (Bergmann and Sack, 2007; Pillitteri and Torii, 2012). An MMC undergoes an asymmetric entry division to generate a meristemoid (M) and a sister stomatal lineage ground cell (SLGC). A SLGC can either differentiate into a pavement cell or undergo asymmetric spacing division to produce a meristemoid and another SLGC. A meristemoid undergoes several rounds of asymmetric amplifying divisions before finally differentiating into a guard mother cell (GMC), which then undergoes a symmetric division to produce two terminal young guard cells (GCs). The young GC pair then undergo a maturation process to become a functional mature stoma.

Many key regulators of early stomatal development that leads to the production of GMCs and young GCs in Arabidopsis have been identified in the past two decades (Bergmann and Sack, 2007; Pillitteri and Torii, 2012; Zoulias et al., 2018). In contrast, much less has been described about how young GCs develop into functional mature stomata. Two known regulators of GC maturation are FAMA, a bHLH transcription factor (TF) critical for GC maturation and fate maintenance, and SCAP1, a Dof-like TF regulating mature GC morphologies and cell wall lining (Matos et al., 2014; Negi et al., 2013). Compared to other cells, cell walls of mature GCs are modified to have properties like extra strength, enhanced flexibility, and anisotropy (Rui et al., 2018; Yi et al., 2019). Among them, anisotropy describes the phenomenon that a material has direction-dependent response to applied force. In GC cell walls, anisotropy is imparted by circumferentially-oriented cross-linked cellulose microfibrils (CMFs), which provide GCs with hoop rigidity (Woolfenden et al., 2017). Hoop rigidity constrains GCs from enlarging their widths in the cross-section direction upon increased turgor pressure, which is crucial for GC movement, especially openings (Aylor et al., 1973; Woolfenden *et al*., 2017). However, the key regulators of GC cell wall properties remain largely uncharacterized, although some related modifying enzymes have been described, such as PME6, PMEI18, PGX3, PLL12, GAUT10, and GAUT11 (Amsbury et al., 2016; Chen et al., 2021; Guo et al., 2021; Rui *et al*., 2018; Zhang et al., 2023).

Another important aspect of stomatal physiology is, once they are mature, what are the regulators of their movement in response to different abiotic and biotic conditions. A number of signaling, kinase, and transporter genes participate in the process, like the ABA singling genes *ABI1* and *OST1*, kinases genes *MPK9* and *MPK12*, and transporter genes *SLAC1*, *QUAC1*, and *GORK* (Allen et al., 1999; Hosy et al., 2003; Jammes et al., 2009; Merlot et al., 2002; Meyer et al., 2010; Mustilli et al., 2002; Negi et al., 2008; Vahisalu et al., 2008). Some of these genes show preferential expression in mature GCs, such as *MPK9*, *MPK12*, *SLAC1* and *QUAC1*. However, due to the complex signaling pathways and transporter systems that modulate stomatal movement, there might be more relevant regulators remain to be identified. A complete catalogue of the genes specifically expressed in mature GCs might help to identify such unknown regulators.

A number of studies based on bulk-transcriptome analysis have analyzed transcriptomes of mature GCs previously. For example, purified GC protoplasts have been used to profile mature GC transcriptomes and isolated a GC-specific gene promoter *pGC1* (Bates et al., 2012; Leonhardt et al., 2004; Yang et al., 2008). In another study, stomatal lineage cells at different developmental stages were labelled with stage-specific fluorescent proteins, and stage-specific cells were isolated by fluorescence-activated cell sorting and used for transcriptome profiling by RNA-seq, from which gene clusters with stage-specific expression patterns were identified (Adrian et al., 2015). Recently, with the advent of single-cell technologies, single-cell RNA-sequencing (scRNA-seq) analyses of enriched GCs, cotyledons, or apices of Arabidopsis have also been reported, from which stomatal lineage cells at different developmental stages were identified (Liu et al., 2020b; Lopez-Anido et al., 2021; Zhang et al., 2021). Using these datasets, it become possible to identify genes with stage-specific expression during GC development. However, due to the intrinsic noise and data sparsity contained within scRNA-seq data, analyzing a single dataset alone might retrieve genes with noised expression, and integration of multiple datasets can help to obtain a more reliable list of GC-specific genes to guide down-stream gene function studies.

In the current study, we developed a data analysis scheme to enable cross-dataset integration of stomata-related scRNA-seq datasets. We used the scheme to conduct a maturomics analysis of GCs to identify genes regulating GC maturation and function formation. We used the term “maturomics” to denote omics analysis of the maturation and function formation process of a tissue or organ. After integrating three publicly available scRNA-seq datasets, we obtained a list of 586 genes that are specifically up-regulated during GC maturation. The list is enriched with regulators of GC functions. From the gene list, we identified and verified a pair of G2-like TF genes as regulators of stomatal size and GC hoop rigidity, whose double mutants had severe defects in regulating stomatal pore apertures. Our analysis thus provides a reliable gene list for studying GC maturation and function formation.

## Results

### Identification of guard cells of different developmental stages across single-cell datasets

We aimed to identify genes regulating stomatal maturation and function formation using single-cell transcriptome datasets. To assist single-cell data analysis, we first used an Arabidopsis gene co-expression network, AtGGM2014, to extract gene co-expression modules related to stomata and epidermis. AtGGM2014 is a gene co-expression network we constructed previously using large-scaled microarray-based bulk transcriptomes, from which 662 gene co-expression modules functioning in diverse biological processes were identified (Ma et al., 2015). Among them, Module #76 and #118 are enriched with genes possessing the gene ontology (GO) terms *stomatal movement* and *stomatal complex development*, respectively (BH-adjusted *P*-value, *P* = 2.32E-06 and 9.20E-19) (**Figure 1A, B, and Table S1**). #76 contains 53 genes in total, and at least 20 of them are known regulators of guard cell (GC) maturation or functions, such as *SCAP1* and *OSP1* that regulate GC maturation and structural formation, as well as *SLAC1*, *BLUS1*, and *MPK12* that are expressed in mature GCs and regulate GC movement (Jammes *et al*., 2009; Negi *et al*., 2008; Negi *et al*., 2013; Takemiya et al., 2013; Tang et al., 2020; Vahisalu *et al*., 2008). We thus considered #76 as a gold standard gene set regulating GC maturation and functions and used it as a marker module for GCs. In contrast, #118 contains stomatal development regulators expressed in meristemoid mother cells (MMCs), meristemoids (Ms), or stomatal-lineage ground cells (SLGCs), such as *SPCH*, *BASL*, and *TMM*, or in guard mother cells (GMCs), such as *MUTE* (Dong et al., 2009; MacAlister et al., 2007; Nadeau and Sack, 2002; Pillitteri et al., 2007; Yang and Sack, 1995). We considered #118 as a marker module for early stomatal development. Similarly, #37 is a marker module for epidermal cells, containing epidermal cell marker genes *ATML1*, *PDF1*, and *FDH*, and enriched with genes involved in the *wax biosynthetic process* (*P* = 3.43E-14); and #8 is a marker module for cell cycle, which is enriched with 26 genes for *cytokinesis* (*P* = 1.10E-21), such as *AUR3*, *POK1*, *POK2*, and *RUK* (**Figure 1C, D and Table S1**).

**Figure 1.**
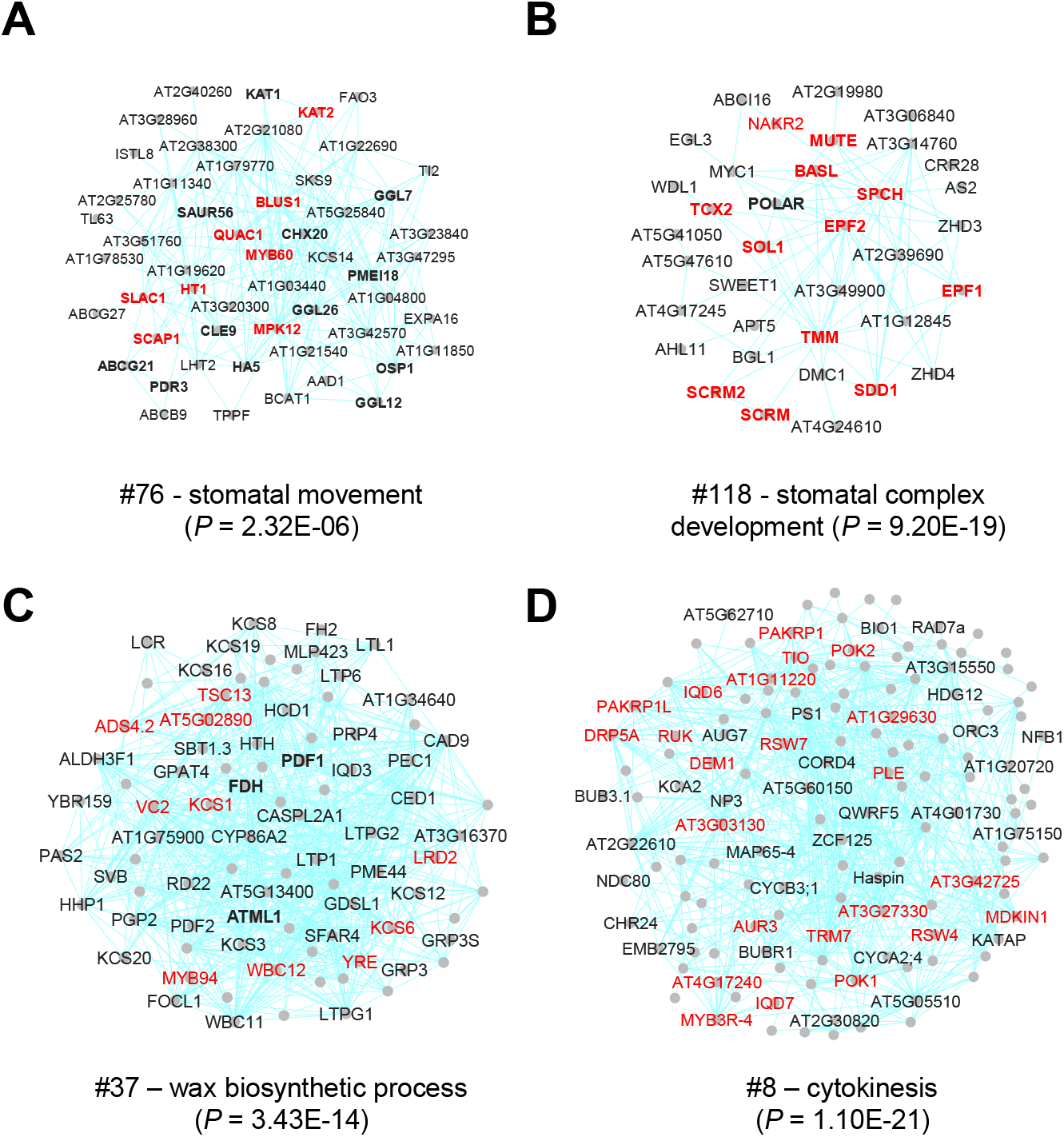
Stomata and epidermis-related gene co-expression modules. The modules are extracted from an Arabidopsis gene co-expression network AtGGM2014. **(A)** Module #76 involved in stomatal maturation and functions. Node represent genes, and connections between genes indicate they have co-expression relationships. Genes in red text indicated they have the GO term listed below the network. Same as below. Genes highlighted in bold are known regulators of stomatal maturation or functions (see Table S1 for a complete reference). **(B)** Module #118 involved in stomatal development. Genes highlighted in bold are known regulators of stomatal development. **(C)** Module #37 involved in epidermal functions. Genes highlighted in bold are markers of epidermis cells. **(D)** Module #8 involved in cytokinesis.

We used the modules to extract and analyze epidermal and stomatal cells from three Arabidopsis scRNA-seq datasets published previously. The first dataset, called “SHOOT1” here, was obtained from Zhang et al. and contains cells sampled from shoot apices of 7-day-old Arabidopsis seedlings (Zhang *et al*., 2021). A clustering analysis identified 40 cell clusters from the dataset, among which 12 were considered as epidermal or stomatal cells due to their expression of Module #37 (**Fig. 2A and S1A**). The cells in Clusters 31, 30, 10, and 33, expressing Module #118 and *SPCH*, were considered as early stomatal lineage cells like MMCs, Ms, and SLGCs; a subset of the cells in Cluster 35, expressing both #118 and *MUTE*, were labeled as GMCs, another group of early stomatal lineage cells; those in Cluster 17 and a small subset of cells in Cluster 35, expressing Module #76 and *FAMA*, were labeled as GCs; those in Clusters 15, 32, 27 and 23 were labeled as epidermal cells due to their expression of Module #37 and *ATML1* (**Figure 2B**); while other cell clusters not expressing Module #37 were considered as non-epidermis (**Fig. S1B**). Interestingly, Module #8, an indicator of cytokinesis activities, is expressed in three clustered areas I – III, which largely overlap with Cluster 33, 10, and 35 (partial) (**Figure 2C**). In early stomatal development, SLGCs and Ms undergo two types of asymmetric cell divisions, including spacing and amplifying divisions, to generate more Ms and pavement cells, and *SPCH* is expressed along the processes (MacAlister *et al*., 2007). At a later time-point, each GMC undergoes one round of symmetric cell division to produce a pair of terminal young GCs, which is regulated by MUTE and FAMA, and from that point the GC maturation process starts (Ohashi-Ito and Bergmann, 2006; Pillitteri *et al*., 2007). Based on the expression of relevant genes, we considered that the cells in areas I and II are SLGCs and Ms undergoing asymmetric division, while those in area III are GMCs undergoing symmetric division (**Figure 2C**).

**Figure 2.**
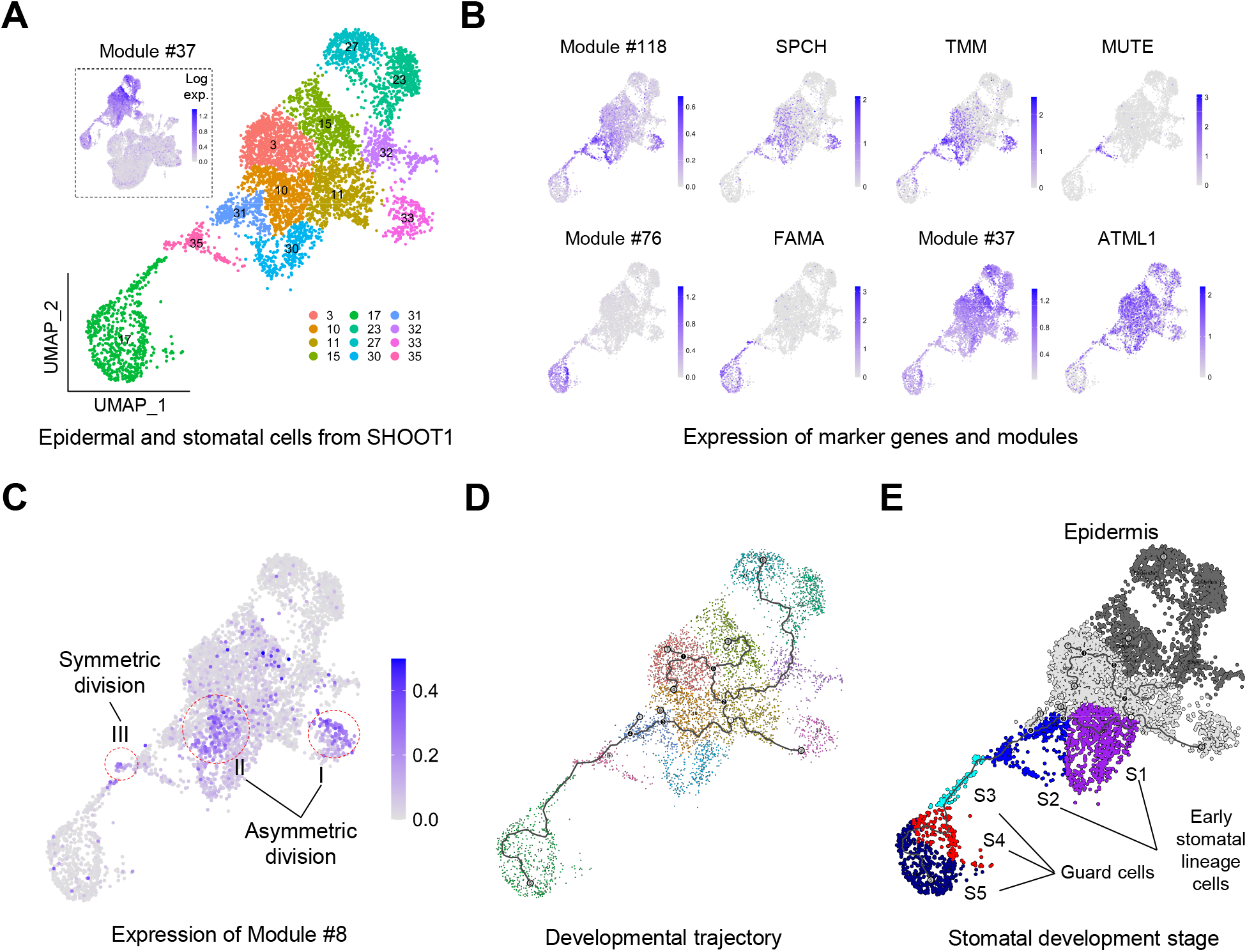
Identification of stomatal lineage cells at different developmental stages from the SHOOT1 dataset. The dataset is obtained from Zhang et al. (2021). **(A)** Extraction of epidermal and stomatal cells from SHOOT1. The cells from SHOOT1 were clustered into 40 clusters (see Figure S1A for a complete UMAP plot). The box on the top left shows the expression level of Module #37 among all cells. The cell clusters expressing Module #37 were extracted as epidermal and stomatal cells, including clusters 3, 10, 11, 15, 17, 23, 27, 30, 31, 32, 33, 35. **(B)** Expression levels of Module #37, #76, #118, and marker genes *SPCH*, *TMM*, *MUTE*, *FAMA*, and *ATML1* in the epidermal and stomatal cells. **(C)** Expression level of Module #8. Three areas with clustered expression are highlighted. Two areas are considered to have cells undergoing asymmetric divisions (I and II), while another area is considered to have cells undergoing symmetric division (III). **(D)** The cells’ developmental trajectories. **(E)** Five developmental stages assigned to stomatal lineage cells according to their developmental trajectories. Epidermis cells are highlighted as a control. The early stomatal lineage cells include mother meristemoid cells (MMCs), meristemoinds (Ms), stomatal lineage ground cells (SLGCs), and guard mother cells (GMCs).

We then inferred developmental trajectories for both stomatal and epidermal cells using Monocle 3 and divided the stomatal trajectory into five different stages (Trapnell et al., 2014) (**Figure 2D, E**). To enable cross-dataset comparison, we used key developmental events to calibrate stomatal developmental stages along the stomatal trajectory between different scRNA-seq datasets. The starting point of stomatal trajectory was chosen just outside the cell clusters expressing Module #118, which can be considered as the junction between stomatal lineage cells and epidermal cells. The cells that express both *FAMA* and Module #8 where symmetric division is supposed to occur, which are also positioned right after the cells expressing *MUTE*, were considered as the starting point of GC maturation. The section of the trajectory before the starting point of maturation was divided equally into stages S1 and S2, while the section after was divided equally into stages S3, S4, and S5 (**Figure 2E**). Thus, S1 and S2 represent the early stages of stomatal development, covering early stomatal lineage cells, including MMCs, Ms, SLGCs, and GMCs, while S3, S4, and S5 represent the stages of GC maturation and function formation. By using S1 and S2 cells and non-epidermal cells as controls, we can investigate gene expression dynamics in S3 to S5 to identify genes regulating stomatal maturation and function formation.

Following a similar strategy, stomatal and epidermal cells were also identified from another two stomata-related scRNA-seq datasets. One dataset, named as “ATML” here, was obtained from Lopez-Anido et al., where the authors conducted scRNA-seq for stomata via enriching epidermal cells by sorting epidermal cells labeled by YFP under the control of *ATML1* promoter (**Figure 3A-C and S1C, D**) (Lopez-Anido *et al*., 2021). The other dataset, named as “COTYLEDON” here, was obtained from Liu et al., where five-day-old cotyledons, enriched with stomatal cells, were analyzed by scRNA-seq (**Figure 3D-F and S1E, F**) (Liu *et al*., 2020b). In ATML, Module #118 and #76 also helped to identify early stomatal lineage cells and GCs, respectively. Module #8 is also expressed in the cells between early stomatal lineage cells and GCs, where FAMA also starts expressing. However, compared to the SHOOT1 dataset, less cells are expressed Module #8, which could be due to less young stomatal lineage cells were capture in this sample. But this should not affect the identification of five stages of stomatal development using the same method as above (**Figure 3C**). In the COTYLEDON dataset, even less early stomatal lineage cells expressing #118 were identified, but we can still use the expression of *FAMA* to mark the cells at the starting of GC maturation (**Figure 3E**). Additionally, the number of epidermal cells expressing Module #37 but not GC markers are also scarce, and we therefore chose the cells at the boarder of the cell cluster expressing #118 as the starting point of GC developmental trajectory, and five developmental stages were also identified accordingly (**Figure 3F**).

**Figure 3.**
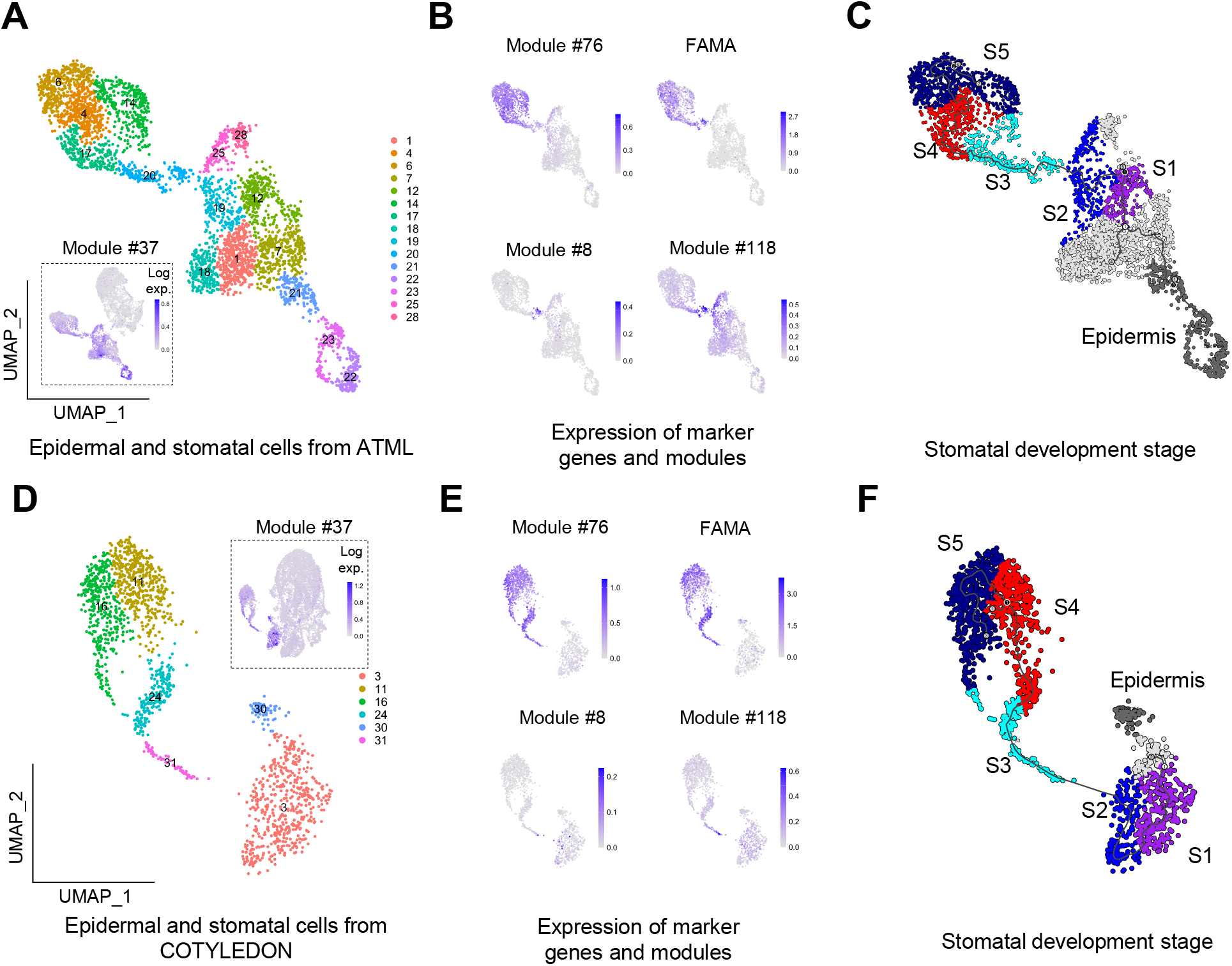
Identification of stomatal lineage cells at different developmental stages from the ATML and COTYLEDON datasets. These two datasets are obtained from Lopez-Andido et al. (2021) and Liu et al. (2020a). **(A, D)** Extraction of epidermal and stomatal cells from ATML and COTYLEDON. The complete UMAP plots for the two datasets are shown in Figure S1. **(B, E)** Expression levels of Module #8, #76, #118 and *FAMA* in the extracted cells. **(C, F)** Developmental stages assigned to stomatal lineage cells in the ATML and COTYLEDON datasets.

### Genes up-regulated during guard cell maturation

After obtaining GCs at different developmental stages, we used two differential gene expression analyses to identify genes up-regulated during GC maturation (**Figure 4A**). Firstly, we identified 964, 3979, and 942 significantly up-regulated genes (FC ≥3 and P-value < 0.01) in GCs (S3, S4, and S5 combined) compared to non-epidermis cells in the SHOOT1, ATML, and COTYLEODON datasets, respectively; among them, 450 genes were commonly up-regulated in all of these three datasets. Secondly, we also identified 1387, 1567, and 814 significantly up-regulated genes in GCs compared to early stomatal lineage cells (including MMCs, Ms, SLGCs, and GMCs; S1 and S2 combined) in the three scRNA-seq datasets respectively, and 350 genes were commonly up-regulated in all datasets (**Figure 4B and Table S2**). The union of these two analyses identified a list of 586 genes commonly up-regulated in GCs in all three datasets. We considered this gene list as a core collection of genes up-regulated during stomatal maturation and named the list as sc_586 (**Table S3**). Selected genes from sc_586 were plotted for their expression levels along stomatal developmental stages in the three datasets, which clearly demonstrated obvious trends of up-regulation during stomatal maturation (**Figure 4C**).

**Figure 4.**
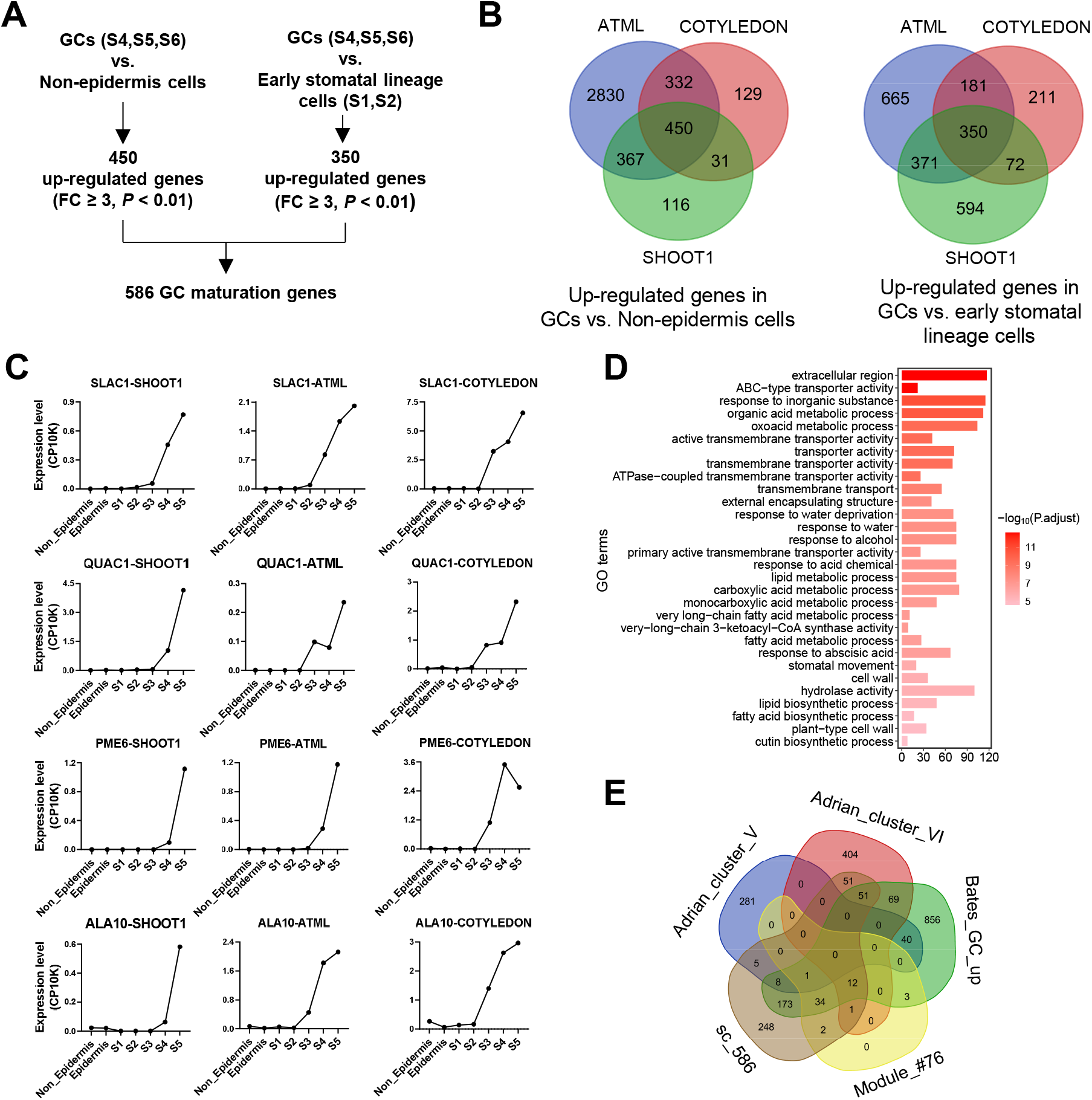
Identification of sc_586, a gene list specifically up-regulated during stomatal maturation and function formation. (**A**) Two differential gene expression analyses were used to identify 586 genes specifically up-regulated during stomatal maturation and function formation. (**B**) Venn diagrams showing the genes commonly up-regulated in all three scRNA-seq datasets. (**C**) Stage-specific expression of genes selected from sc_586 in three scRNA-seq datasets. **(D)** Enriched GO terms from sc_586. **(E)** A Venn diagram comparing mature stomata-related gene lists. Among them, Module #76, extracted from AtGGM2014 gene co-expression network, contains 53 genes and is considered as a gold standard gene set for stomatal maturation and function formation; sc_586 is the list identified from this study; Bates_GC_up is a gene list identified to be up-regulated in GCs compared to leaves identified by Bates et al. (2012); and Adrian_Cluster_V and Adrian_Cluster_VI are two gene modules up-regulated in mature stomata identified by Adrian et al. (2015).

A GO analysis indicated that sc_586 is enriched with genes involved in *stomatal movement* or with *transporter activity* (*P* = 8.45E-07 and 3.26E-10) (**Figure 4D**). The list contains at least 69 known stomatal genes, most of which regulate stomatal maturation or functions (**Table 1 and Table S3**). 20 of these known regulators are among the known regulators contained within Module #76, as described in the previous section (**Figure 1A**). Additionally, sc_586 contains another 49 known stomatal regulators not present in Module #76, such as transporter genes *GORK*, *CLC-A*, *AHA1*; TF genes *FMA*, *SCRM*; kinase genes *NUT*, *RAF6*, *MPK9*; and enzyme genes *GPAT4*, *GPAT8*, *PME6* (Amsbury *et al*., 2016; Fernández-Santos et al., 2020; Hosy *et al*., 2003; Hsu et al., 2021; Jammes *et al*., 2009; Kanaoka et al., 2008; Li et al., 2007; Liu et al., 2020a; Liu et al., 2022b; Matos *et al*., 2014; Ohashi-Ito and Bergmann, 2006; Peng et al., 2022; Wege et al., 2014; Yamauchi et al., 2016). The list also contains un-characterized genes encoding transporters, TFs, kinases, enzymes, and other types of proteins (**Table 1 and Table S3**). As these genes were specifically up-regulated during GC maturation, they could be considered as ideal candidate genes for future studies on stomatal maturation and functions.

**Table 1.**
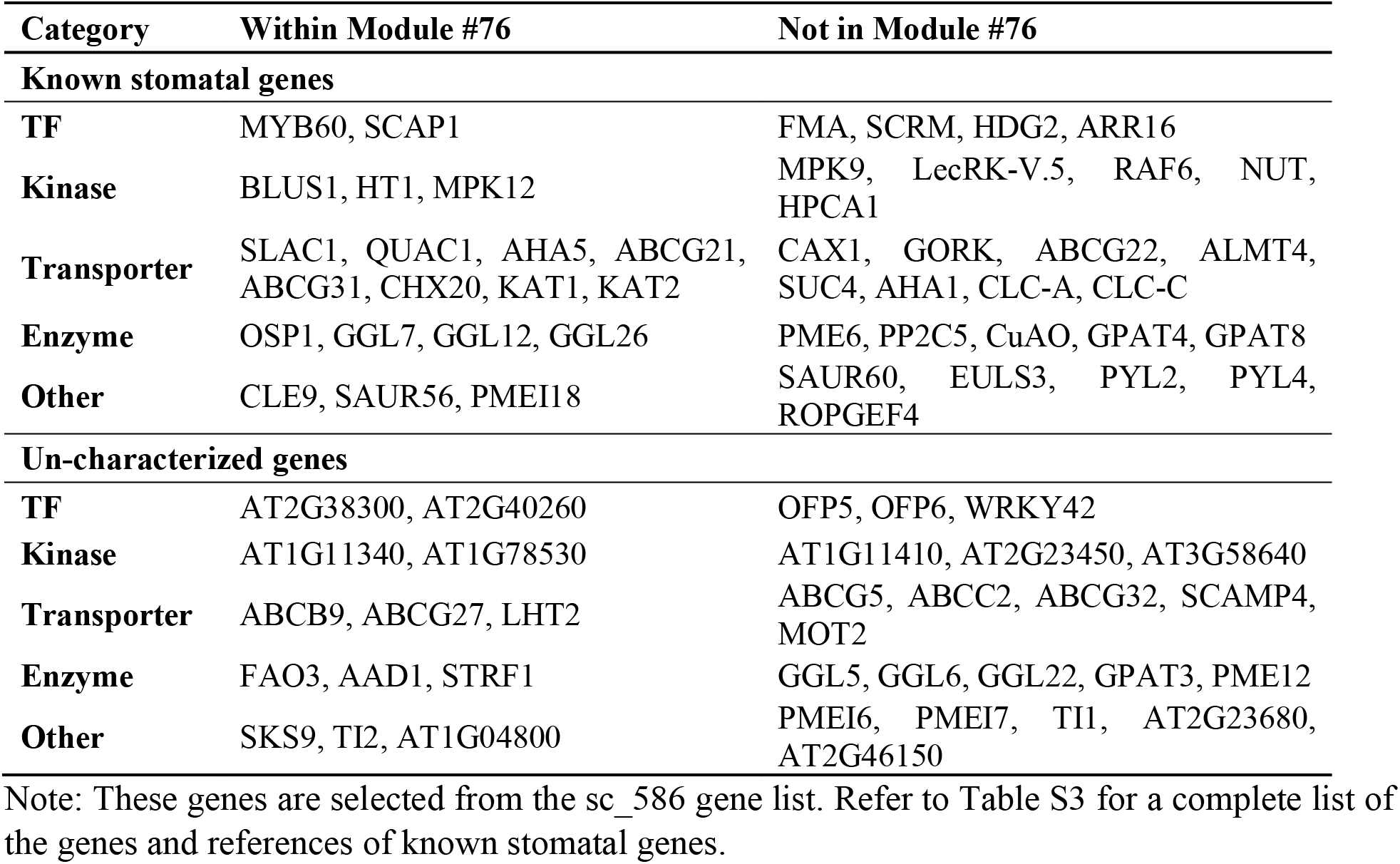
Genes up-regulated during stomatal maturation and function formation (partial)

We noted that previous studies based on bulk transcriptome analysis have also identified genes highly expressed in GCs (Adrian *et al*., 2015; Bates *et al*., 2012; Leonhardt *et al*., 2004; Yang *et al*., 2008). Among the two most recent studies, Bates et al. identified a list of 1247 up-regulated genes (named as Bates_GC_up here) in GCs compared to leaves; Adrian et al. measured the transcriptomes of stomatal lineage cells using RNA-seq and identified two gene clusters (named as Adrian_ClusterV and Adrian_ClusterVI here) specifically expressed in mature stomata that contains 335 and 588 genes respectively (Adrian *et al*., 2015; Bates *et al*., 2012). We extracted these genes lists and, together with sc_586, compared their overlaps with the genes in Module #76, which contains 53 genes and serves as an independent gold standard for genes regulating GC maturation and functions (**Figure 4E and Table S4**). While Adrian_ClusterV and Adrian_ClusterVI have only 1 and 12 genes overlapped with Module #76, sc_586 and Bates_GC_up both have 50 genes overlapped with Module #76. The result indicated that sc_586 and Bates_GC_up have higher capabilities than Adrian_ClusterV and Adrian_ClusterVI to recapitulate genes involved in stomatal maturation and functions. Additionally, since sc_586 contains 586 genes and Bates_GC_up contains 1237 genes and they both recapitulate the same number of genes from Module #76, the genes in sc_586 could be more relevant to stomatal maturation and functions than those in Bates_GC_up. In other words, sc_586 serves as a better source to retrieve candidate genes for reverse genetics studies on stomatal maturation and functions.

### Expression of gene families during stomal development

Besides identifying genes involved in stomatal maturation and functions, our analysis also enables visualizing expression patterns of gene families during stomatal development across scRNA-seq datasets. Due to functional redundancy, such visualization can be helpful to pinpoint relevant candidate genes for functional studies. As an example, the MAPK kinase gene family plays important roles in responses to environmental signals, including regulating stomatal movement. The Arabidopsis genome contains 20 MAPK genes. These 20 genes showed a wide range of expression patterns during stomatal development (**Figure 5A**). Among them, *MPK9* and *MPK12* are contained within sc_586 and they showed clear up-regulation during stomatal maturation (stages S3 – S5) in all three scRNA-seq datasets. Other MPK genes also had expression in GCs, such as *MPK4*, *MPK17*, *MPK6*, and *MPK19*, although their expressions were not as specific as *MPK9* and *MPK12*. As another example, we also visualized the expression of the G2-like TF gene family (**Figure 5B**). This family contains 40 genes in total, and a number of them showed expressions in guard cells in all three scRNA-seq datasets, such as *HHO2*, *HHO3*, *GammaMYB2*, *AT2G38300*, and *AT2G40260*. However, among these genes, only *AT2G38300* and *AT2G40260* showed mature guard cell specific expression, and these two genes are also contained within our sc_586 gene list, while others like *HHO2*, *HHO3* and *GammaMYB2* also had expression in epidermis cells. Interestingly, among all G2-like TFs, AT2G38300 and AT2G40260 are most close to each other phylogenetically (**Figure 6A**). The next closely related G2-like TF, AT2G42660, does not have expression in GCs (**Figure 5B**). We thus hypothesized that *AT2G38300* and *AT2G40260* might play roles in stomatal maturation or function formation.

**Figure 5.**
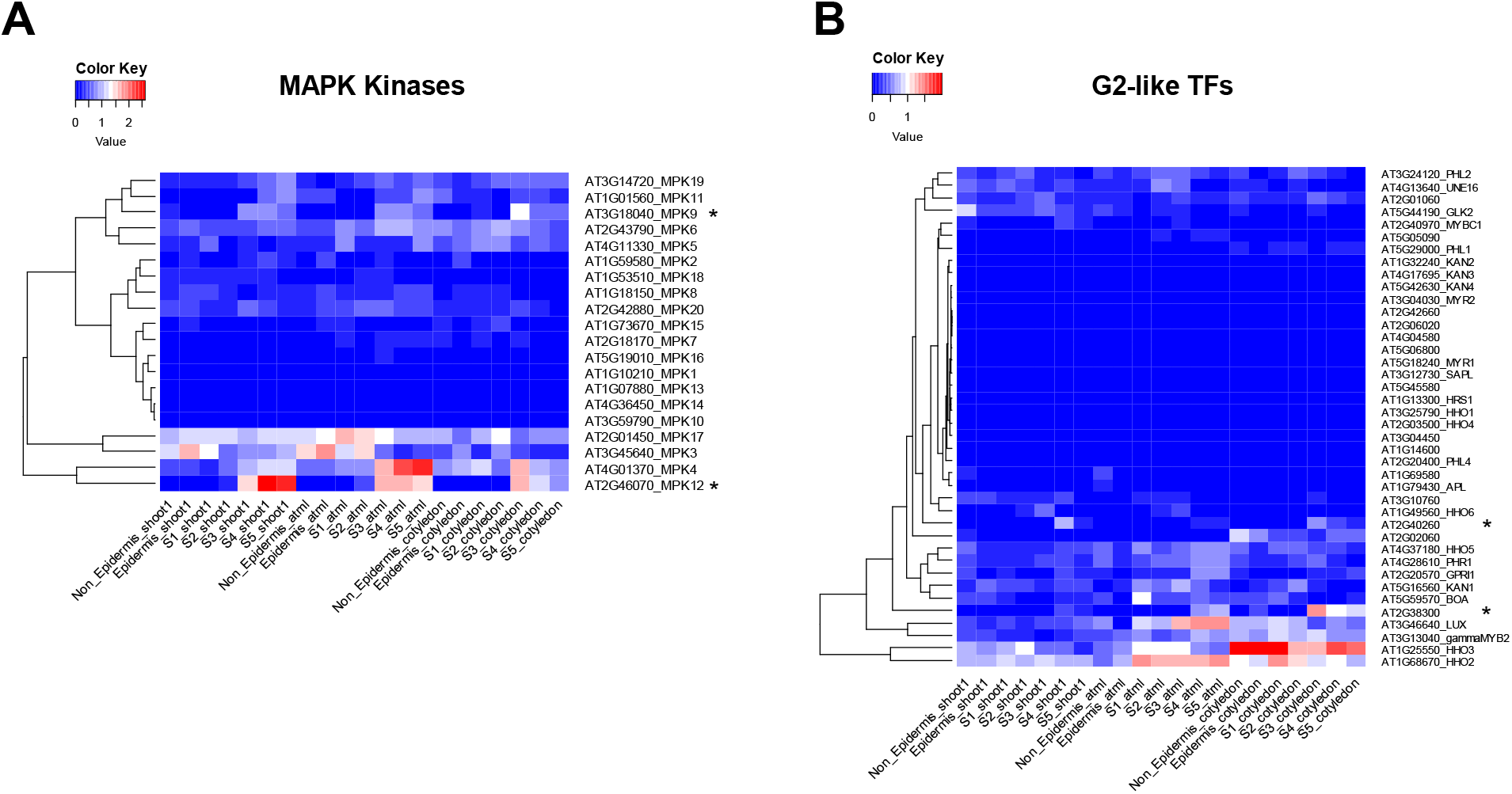
Two heatmaps showing stage-specific expression of MAPK (A) and G2-like TF (B) gene families during stomatal development and maturation across three scRNA-seq datasets. Values are log_2_(CP10K) gene expression values. “*” marks the genes contained within sc_586.

**Figure 6.**
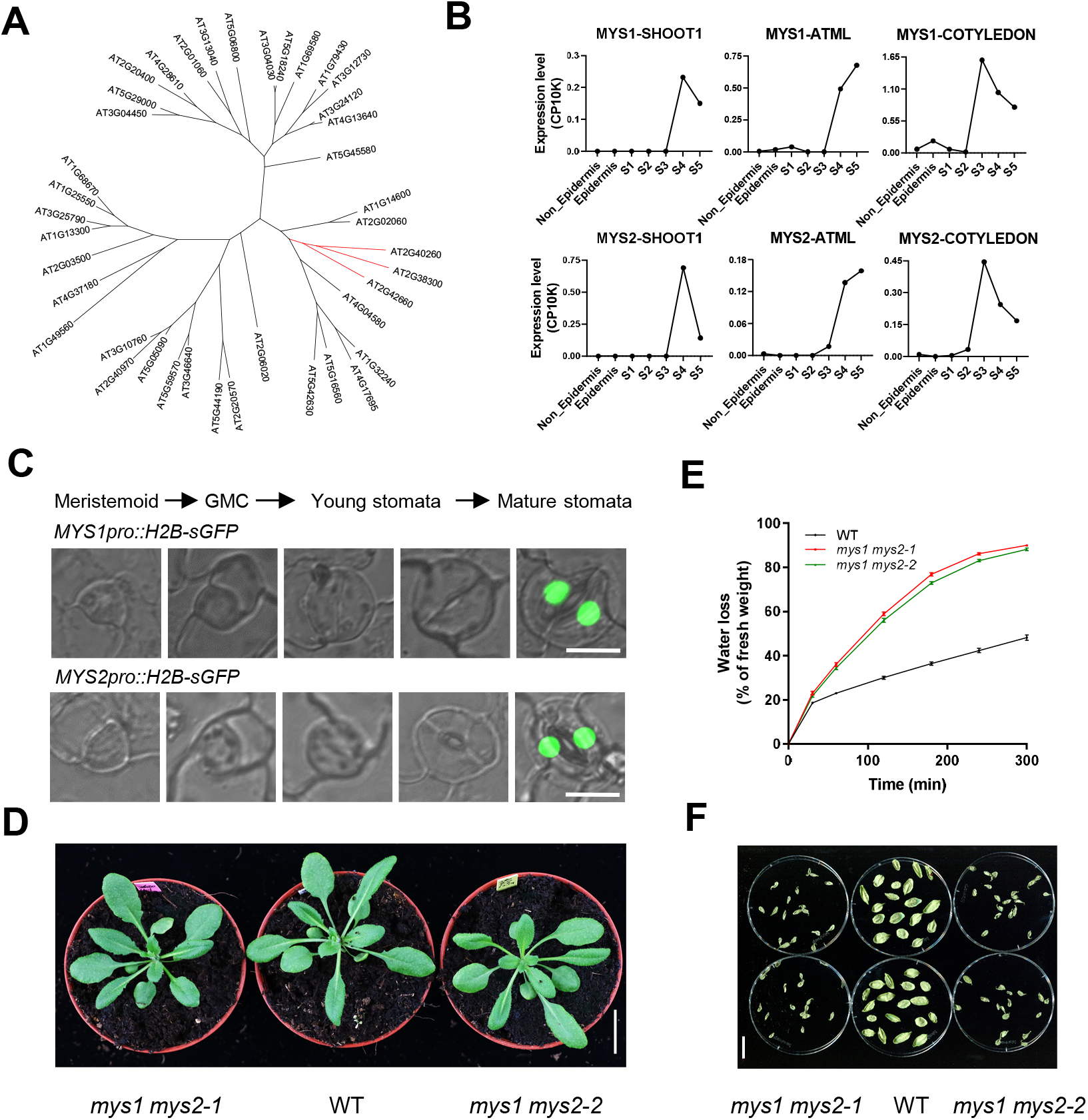
Expression patterns of *MYS1* and *MYS2* and phenotypes of their double mutants. (**A**) A phylogenetic tree for all G2-like TF proteins in Arabidopsis. The phylogenetic tree was constructed using the neighbor-joining method in the MEGA6 software (Tamura et al., 2013). **(B)** Expression of *MYS1* and *MYS2* in the three scRNA-seq datasets. **(C)** Expression patterns of *MYS1* and *MYS2* in stomatal lineage cells. Bar, 10 μm. **(D)** Mutant lines of *mys1 mys2* under normal growth condition. Bar, 2 cm. **(E)** Water loss of detached leaves of *mys1 mys2-1*, *mys1 mys2-2*, and WT plants. Values are means ± SE (n=3). **(F)** Leaves of *mys1 mys2-1*, *mys1 mys2-2*, and WT plants 5 hours after detachment. Bar, 2 cm.

### Verification of two G2-like TFs as regulators of stomatal size and guard cell hoop rigidity

We tested if *AT2G38300* and *AT2G40260* regulate stomatal maturation or functions. Of note, in a recent study these two genes have been named as *MYS1* (*MYB-SHAQKYF 1*, *AT2G38300*) and *MYS2* (*AT2G40260*) (Liu et al., 2022a). The authors found that *MYS1* and *MYS2* were co-expressed with wax biosynthesis genes *GGL7* (*AT1G33811*), *GGL26* (*AT5G18430*), *OSP1* (*AT2G04570*), and *CER26-Like* (*AT3G23840*), and they hypothesized and verified that *MYS1* and *MYS2* are regulators of wax biosynthesis and drought tolerance. Interestingly, all these wax biosynthesis genes are also contained within our sc_586 gene list as well as within Module #76 of AtGGM2014 (**Figure 1A and Table S3**), indicating they all have GCs specific expression patterns. Although the previous study also found by using GUS staining that *MYS1* and *MYS2* are predominantly expressed in GCs (Liu *et al*., 2022a), their roles in GC development and function formation have not been fully addressed. Here, we kept their gene names and asked whether these two genes also participate in GC maturation and function formation.

*MYS1* and *MYS2* displayed mature GCs-specific expression patterns in all three scRNA-seq datasets (**Figure 6B**). To examine their cell-type specific expression patterns more closely, we generated Arabidopsis transgenic lines expressing the fusion protein H2B-sGFP driven by the *MYS1* and *MYS2* promoters. Both lines showed that *MYS1* and *MYS2* are specifically expressed in mature GCs (**Figure 6C**). We noted the expression pattern of *MYS1* is similar to that has been reported before (Zhang *et al*., 2021). Additionally, transiently expressed MYS1-GFP and MYS2-GFP fusion proteins driven by the 35S promoters in *Nicotiana benthamiana* leaves mainly localized to the nucleus, supporting their roles as TFs (**Figure S2**), which is also consistent with the results by Liu et al. (2022a).

To investigate their physiological functions, we generated single mutants of *MYS1* and *MYS2* via the CRISPR technology, by using two sgRNAs, sgRNA-300 and sgRNA-260, to target *MYS1* and *MYS2* respectively (**Figure S3A, B**). The mutations in these two single mutant lines led to early termination of MYS1 and MYS2 proteins biosynthesis respectively (**Figure S4A**). However, these two single mutants did not display obvious phenotypes and their leaves lost water at the same rates as WT plants (**Figure S4B**). Considering their close homology, we generated their double mutants by using a single CRISPR construct containing both sgRNA-300 and sgRNA-260 **(Figure S3B**). Two independent double mutant lines, *mys1 mys2-1* and *mys1 mys2-2*, were obtained (**Figure S5A, B**). In *mys1 mys2-1*, *MYS1* had 1 bp insertion and *MYS2* had 2 bp insertion plus 12 bp deletion near the sgRNA sites, resulting in early termination of both proteins’ biosynthesis. In *mys1 mys2-2*, *MYS1* had 10 bp insertion and 27 bp deletion and *MYS2* had 53 bp deletion near the sgRNA sites, also resulting in early termination of both proteins’ biosynthesis. Both *mys1 mys2-1* and *mys1 mys2-2* grew normally under normal conditions (**Figure 6D**). However, in detached leaf experiments, both mutant lines lost water at much faster rates than WTs (**Figure 6E**), which is similar to the results obtained by Liu et al (2022a). At 5 hours post detachment, the leaves of *mys1 mys2-1* and *mys1 mys2-2* lost 88% – 90% of their fresh weight and wilted almost completely, while those of WT plants lost 48% of their fresh weight and still maintained their shapes (**Figure 6F**). To further confirm that the mutations of both *MYS1* and *MYS2* caused the water loss phenotypes, we generated another two independent double mutant lines *mys1 mys2-3* and *mys1 mys2-4* via CRISPR by using a second set of sgRNAs, sgRNA-300-2 and sgRNA-260-2, whose sequences were different from sgRNA-300 and sgRNA-260 (**Figures S3A, B and S6A, B**). In *mys1 mys2-3*, *MYS1* has 1 bp insertion and *MYS2* has another 1 bp insertion near the sgRNA sites, and in *mys1 mys2-4*, *MYS1* has 22 bp deletion and *MYS2* has 4 bp insertion near the sgRNA sites, all leading to early termination of the corresponding proteins’ biosynthesis. These two independent double mutant lines also lost water significantly faster than WTs (**Figure S6C**).

We then examined stomatal morphologies of the double mutants in details. Leaves from WT and *mys1 mys2* mutant lines were immersed in stomatal opening buffer under light and epidermal peels were sampled as control (Ctrl) for microscopic observation; the remaining leaves were subjected to abscisic acid (ABA) treatment and then sampled again for observation (**Figure 7A, B**). Stomatal sizes of the double mutants were significantly larger than those of WTs in both control condition and after ABA treatment (**Figure 7C**). Compared to WT, the widths of *mys1 mys2* guard cells were abnormal, resulting in changed stomatal shapes (**Figure 7D**). In WT Arabidopsis, stiffness of GC cell wall is direction dependent, or anisotropic, with the stiffness in the radial directions being reinforced by cross-linked cellulose microfibris (CMFs), which provides the GCs with hoop rigidity (Woolfenden *et al*., 2017). Upon turgor pressure increase, GCs increase their volumes (up to 25%) mainly by extending in the longitudinal direction with limited changes of size in the cross-section direction (Meckel et al., 2007). Such limitation on cross-section size change is crucial for stomatal pore opening (Aylor *et al*., 1973; Woolfenden *et al*., 2017). Due to such limitation, GC width should remain approximately constant for most part of the cell body: the width measured at the central point of GCs, *W_middle_*, should be roughly equal to that measured at the quarter point, *W_quarter_*. In WT plants, the width ratio between the two (*W_middle_* / *W_quarter_*) was 1.03 under control condition, while such ratio increased significantly in both mutant lines to 1.13 (**Figure 7D**). The ratio difference between WT and mutant lines persisted after ABA treatment. The microscopic pictures of the stomata also showed that, in the mutant lines, GC widths were increased in the central part of the cell body in comparison to other parts, resulting in changed stomata shapes (**Figure 7B**). The results provided a first indication that hoop rigidity might be impaired in the mutants.

**Figure 7.**
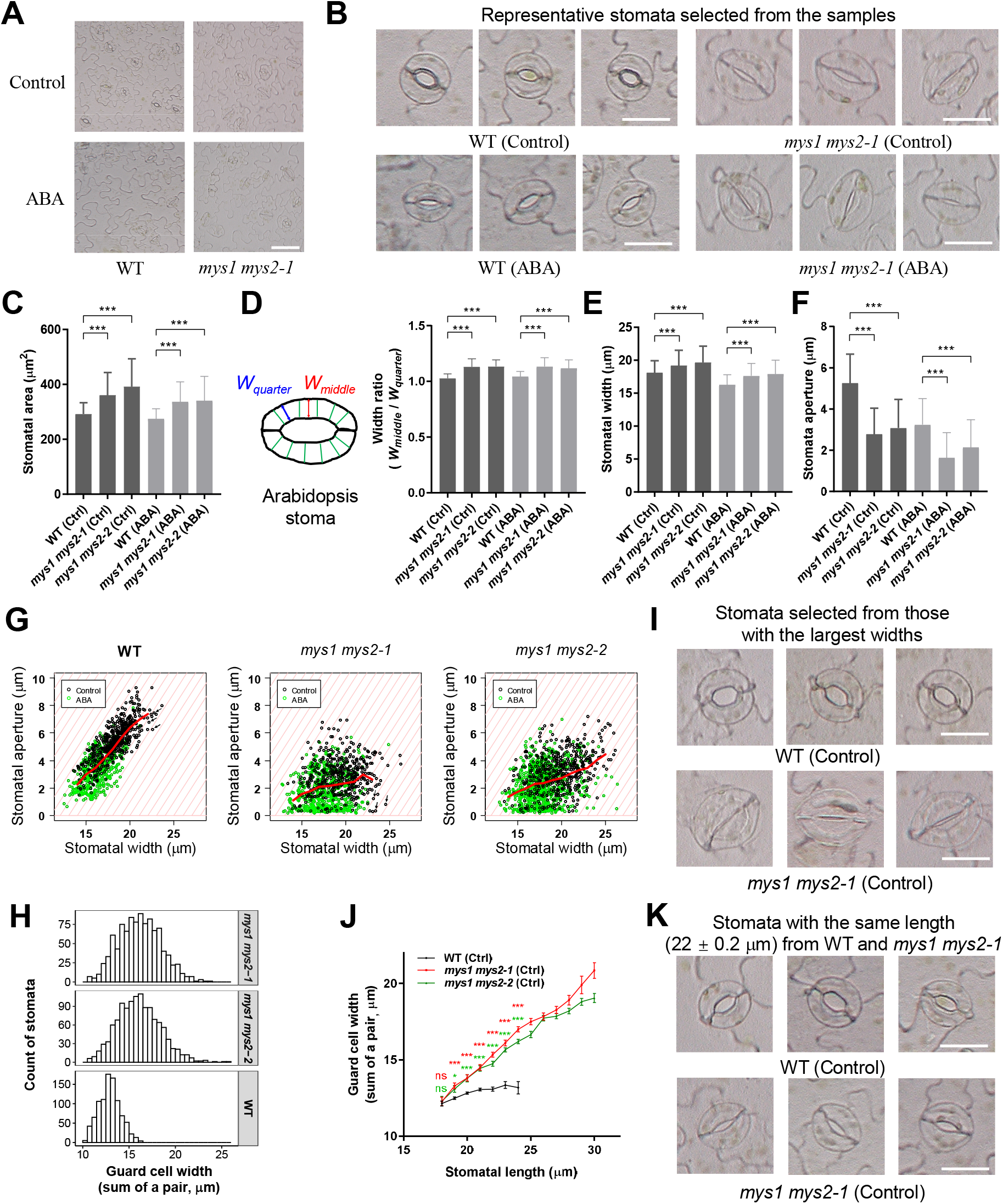
*MYS1* and *MYS2* regulate the size and hoop rigidity of mature guard cells. (**A, B)** Microscopic observations of stomata on epidermis of WT and *mys1 mys2-1* leaves, with (A) showing overviews of the samples and (B) showing enlargement of representative stomata selected from (A). Leaves were immersed in stomata opening buffer under light for 2 hours before epidermal peels were sampled from a portion of the leaves for observation, which were labeled as control samples (Ctrl); the remaining leaves were subjected to 1 μm ABA treatment for another 2 hours before being sampled again, which were labeled as ABA. Samples are the same as below. Bar in (A), 100 μm; Bar in (B), 20 μm. **(C)** Stomatal areas of WT and *mys1 mys2* mutant lines. Values are means ± SD. n = 390 – 708; ***, *P* < 0.001; Student’s t-test. **(D)** Width ratio of guard cells before and after ABA treatment. The diagram on the left shows the definition of *W_middle_* and *W_quarter_*. Green lines indicate the circumferentially-oriented cross-linked cellulose microfibrils that impart GC hoop rigidity. Values are means ± SD. n = 106 – 145; ***, *P* < 0.001; Student’s t-test. **(E, F)** Stomatal widths and stomatal apertures of WT and *mys1 mys2* mutant lines. Values are means ± SD. n = 390 – 708; ***, *P* < 0.001; Student’s t-test. **(G)** Scatter plots of stomata from WT and *mys1 mys2* mutant lines, showing their stomatal widths and stomatal apertures. Each dot represents a stoma from control (green) or ABA-treated (black) samples. Light-red lines in the background represent contour lines showing the difference between stomatal width and stomatal aperture, equal to GC width (sum of a pair). The red lines show the average stomatal apertures at different stomatal widths. (**H**) Histograms showing the distribution of GC widths in the WT and mutant plants. Stomata from both control and ABA-treated samples were combined and used for plotting. **(I)** Selected stomata, indicated by arrows in (G), from WT and *mys1 mys2-1* that have the largest stomatal widths. The stomata with largest widths were first selected from each sample, from which the ones with relatively smaller stomatal apertures were further chosen for displaying. **(J)** A plot showing GC widths of the stomata from WT and *mys1 mys2* mutants against stomatal lengths. Stomatal lengths were rounded to integer numbers before plotting. Only points with ≥ 10 stomata are shown. Values are means ± SE (n = 10 – 140). ns, not significant; *, *P* < 0.05; ***, *P* < 0.001; Student’s t-test. **(K)** Selected stomata with the same length (22 ± 0.2 μm) from WT and *mys1 mys2-1*. Bar, 20 μm.

Additionally, although the double mutant lines had larger stomata than WTs, their stomatal apertures were not enlarged proportionally. The double mutant lines had larger stomatal widths, but their stomatal apertures were significantly smaller than those of the WTs (**Figure 7E, F**). We plotted the widths and apertures of mature stomata from the WT and mutant plants on scatter plots to investigate the relationship between stomatal width and stomatal aperture (**Figure 7G**). The plots used contour lines (light red lines in **Figure 7G**) to illustrate the difference between stomatal width and stomatal aperture, which is equal to the sum of the width of the GC pairs (referred as GC width hereafter). The WT stomata distributed narrowly within a band with GC widths between 10 and 17 μm in the WT plot (**Figure 7G**). The average GC width of WT stomata was 12.91 ± 1.16 μm (mean ± SD), indicating hoop rigidity constrained the expansion or change of GC widths within a small range. However, such constrain on GC width change was reduced or impaired in the mutant lines. The mutant stomata distributed across much larger areas than WT without forming clear bands, and both mutant lines had stomata with GC widths reaching 21 μm and beyond (**Figure 7G**). The average GC widths of the mutant lines were 16.17 ± 2.27 and 16.10 ± 2.30 μm respectively, much larger than those of WTs. The GC widths of the mutants covered larger ranges than that of the WTs (**Figure 7H**), indicating hoop rigidity was reduced or impaired in the mutant lines. As a consequence, stomatal pore movement was severely impaired in the mutant lines. Many stomata in the bottom-right part of the mutant plots had large stomatal widths but small apertures, while similar stomata were not seen in the WT plot (**Figure 7G**). Notably, some stomata with the largest stomatal widths in *mys1 mys2-1* control plants, which had been immersed in stomatal opening buffer for 2 hours under light, still could not open their stomatal pores due to increased GC widths, while WT stomata with similar or even smaller stomatal widths were fully opened (**Figure 7I**), highlighting the importance of GC width constrain in stomatal aperture regulation.

Besides hoop rigidity, another factor that might affect GC widths is stomatal size, as larger stomata tend to have larger GC widths. However, we found that increase in stomatal size alone cannot explain the increase of GC widths in the mutants. We used stomatal length as an indicator of stomatal size, and plotted GC width against stomatal length (**Figure 7J**). In both WT and double mutant plants, larger stomata tend to have larger GC widths; however, the rate of GC width increase vs. stomatal length increase was much faster in the mutant plants. Once stomata length reached 19 μm and beyond, among the stomata with the same lengths, those from mutant plants had significantly larger GC widths than those of WT plants (**Figure 7J**). As an example, selected stomata with the same stomatal length of 22 (± 0.2) μm from WT and mutant plants are shown in **Figure 7K**. Those from the mutant plants had larger GC widths and smaller apertures, and their shapes were apparently different from WTs due to changes in GC width ratio (*W_middle_* / *W_quarter_*). Because mutant stomata of the same size as WTs displayed GC width-related phenotypes, the change of stomatal size alone cannot explain the observed phenotypes. Thus, the results provided another indication that hoop rigidity was impaired or reduced in the double mutant lines. A consequence of such impairment was the double mutants had severe defects in regulating their stomatal pore apertures (**Figure 7E, F, and I**).

Therefore, we concluded that *MYS1* and *MYS2* function redundantly in mature GCs to regulate their size and hoop rigidity. Obtaining hoop rigidity is important for GC maturation and function formation, especially for acquiring the ability to regulate stomatal aperture. The results are consistent with previous modeling studies showing that hoop rigidity is crucial for stomatal pore movement, especially openings (Aylor *et al*., 1973; Woolfenden *et al*., 2017).

## Discussion

In the current study, we integrated three scRNA-seq datasets to identify a gene list sc_586 that are specifically up-regulated during GC maturation and function formation. We used key developmental events to calibrate GC developmental stages between the datasets, so that the GC stages in different datasets can be compared with each other. Our results showed that it is critical to integrate multiple scRNA-seq datasets to obtain a reliable gene list. In the comparison between GCs and non-epidermal cells, 964, 3979, and 942 significantly up-regulated genes were identified from three datasets respectively, but only 450 were common to all three datasets (**Figure 4B**). Results from a single dataset thus could contain a large number of noised genes, while the common ones from all three datasets should reduce such possibility. Indeed, in comparison with other mature GC gene list, our sc_586 showed higher capability to recover the genes in Module #76 of AtGGM2014 that we considered to be a gold standard gene set for GC maturation and function formation (**Figure 4E**). A further inspection of the genes in sc_586 identified many known regulators of GC maturation and functions, indicating other uncharacterized genes with the list could be ideal candidates for future functional studies. Additionally, our analysis also enables visualizing expression of gene families across scRNA-seq datasets, providing additional information to assist candidate genes selection for reverse genetics analysis (**Figure 5 and Table S3**).

As an example, we selected a pair G2-like TF genes, *MYS1* and *MYS2*, from sc_586 for functional verification. A previous study showed that these two genes regulate wax biosynthesis in Arabidopsis leaves and function in drought tolerance response (Liu *et al*., 2022a). However, the study had not fully addressed these two genes’ function in stomata. Our results indicated that *MYS1* and *MYS2* are up-regulated during stomata maturation, and they redundantly regulate the establishment or maintenance of hoop rigidity in mature GCs. Three lines of evidence supported such a conclusion: Firstly, the GC width became unevenly distributed along the cell body in *mys1 mys2* mutant lines. The GC width at the center point of the GC body were significantly enlarged compared to the rest of part, while in WTs GC width stayed approximately equal along the GC body (**Figure 7B, D**). Secondly, GC widths of WT were constrained in a small range among stomata with different stomatal widths, but such constrain was lost or impaired in *mys1 mys2* mutant lines; and some *mys1 mys2* stomata still could not open their pores even their widths had expanded to the largest range after being immersed in stomatal opening buffer for 2 hours (**Figure 7G-I**). Thirdly, for stomata with the same size, *mys1 mys2* stomata had larger GC widths than WTs (**Figure 7J, K**). The results indicated hoop rigidity is reduced or impaired in the mutants, which caused their stomata to have severe defects in stomatal pore aperture regulation. In addition to hoop rigidity, *MYS1* and *MYS2* also regulate stomatal size. It’s possible that the overall cell wall strength and the constrains on GC expansion were reduced in the double mutant lines, as their stomatal sizes were significantly larger than those of the WTs (**Figure 7C**). Thus, *MYS1* and *MYS2* play important roles during GC maturation, which is consistent with their expression patterns during GC development.

In summary, we have identified a gene list sc_586 for GC maturation and function formation from scRNA-seq datasets. We expect this list to facilitate the identification of more regulators of stomatal maturation and functions.

## Materials and methods

### scRNA-seq datasets analysis

The raw data of three public scRNA-seq datasets were downloaded from Beijing Institute of Genomics Data Center and the NCBI SRA database (SHOOT1: samples shoot1_rep1 and shoot1_rep2 of PRJCA003094, http://bigd.big.ac.cn; ATML: SRR13749640; and COTYLEDON: SRR11049211, SRR11049212, SRR11049213, and SRR11049214) as provided by the original studies (Liu *et al*., 2020b; Lopez-Anido *et al*., 2021; Zhang *et al*., 2021). The reads from each sample were combined and mapped to the Arabidopsis genome (araport11) via CellRanger (v5.0.1) to obtain a gene count matrix. Seurat objects were created for each sample using the Seurat (v4.2) package in R (Hao et al., 2021). The cells with ≤200 or ≥5000 expressed genes or with ≥20 percent of the reads mapped to the mitochondria or chloroplast genome were filtered out. The remaining cells were normalized via SCTransform while regressing out the effects of cell-cycle genes, with cell-cycle genes obtained from (Lopez-Anido *et al*., 2021). Principle component analysis (PCA) of the Seurat objects were performed via RunPCA, and the first 50 components were used to conduct further dimension reduction analysis via RunUMAP, with parameters of “min.dist=0.5, dims=1:50”. Cell neighbors were assigned via FindNeighbors, with parameters “reduction=’umap’, dims=1:2, force.recalc=T”, and cell clusters were identified via the FindClusters function.

We then extracted gene co-expression modules (#8, #51, #76, and #118) from the AtGGM2014 gene co-expression network (Ma *et al*., 2015). For each module, we calculated the average expression level of all the genes within the module in every cell, which was considered as the module’ expression level. We then used the expression level of these marker modules as well as known marker genes to identify epidermal, stomatal, and non-epidermal cells from the three scRNA-seq datasets. The epidermal and stomatal cells were then extracted and used for trajectory analysis via Monocle 3 (Trapnell *et al*., 2014). The developmental stages of stomatal lineage cells were assigned accordingly. We then used the FindMarkers function in Seurat to identify differentially expression genes in GCs compared to non-epidermal cells or young stomatal lineage cells, with parameters “logfc.threshold=0, min.pct=0, min.cells.feature=0, pseudocount.use=0.0005”. Genes were deemed as up-regulated in GCs if FoldChange ≥3 and combined P-value < 0.05. We used the Fisher’s method to combine P-values across three scRNA-seq datasets to obtain a combined P-value for each gene. The up-regulated gene list identified by our analysis was compared to the lists identified by others, and Venn diagrams were drawn accordingly using an online tool available at https://bioinformatics.psb.ugent.be/webtools/Venn/.

To obtain developmental stages-specific gene expression values, we first used the raw count values to calculate CP10K values for each gene in each cell. And then for each dataset, we used the cells in each developmental stage to calculate average expression values for every gene in that developmental stage. We removed low expressed genes from further analysis if their CP10K values were smaller than 0.1 in GCs in all three scRNA-seq datasets.

### Plant materials and CRISPR mutant lines

The Col-0 accession of *Arabidopsis thaliana* was used in this study. We used two binary vectors pHY01 and pHY07 developed in our lab (Geng et al., 2023) to generate the CRISPR gene knock-out mutants used in the analysis. Both vectors use a *pNUC-1* gene promoter to drive the expression of *zCas9* gene (Wang et al., 2015) to improve the efficiency of generating CRISPR knock-out mutants. The pHY07 vector is described in (Geng *et al*., 2023), while pHY01 is an earlier version of pHY07. Compared to pHY01, pHY07 is improved in the expression efficiency of mCherry fluorescence proteins used for easy identification of transgenic seeds.

We used CRISP-P (v2.0) to design two sets of sgRNAs to target *MYS1* and *MYS2* (Liu et al., 2017). For *MYS1*, sgRNA-300 (GGTTTCAAGCAATAGTACAG) and sgRNA-300-2 (TTGACCTCCAAGCCTCTCAA) were used; for *MYS2*, sgRNA-260 (AGAAGAATGGAGGAT CCGTG) and sgRNA-260-2 (GCAGCCTCGGAGTCTTTGAG) were used. Single mutants of *mys1-1* and *mys2-1* were generated by using pHY07 vector containing sgRNA-300 or sgRNA-260 only. Double mutants of *mys1 mys2-1* and *mys1 mys2-2* were generated by using a single pHY01 vector containing both sgRNA-300 and sgRNA-260; *mys1 mys2-3* and *mys1 mys2-4* were generated by using a single pHY07 vector containing both sgRNA-300-2 and sgRNA-260-2. The CRISPR vectors were transformed into *Agrobacterium tumefacien* GV3101 and used to transform WT Arabidopsis via the floral dip method (Clough and Bent, 1998). The F1 transgenic plants were sequenced to identify lines containing mutated *MYS1* and/or *MYS2* genes, which were then self-crossed to obtain homozygous mutants. The offspring of the homozygous mutants were then used for further experiments.

### Plant growth conditions, water-loss assays and microscopic observation

The seeds of WT and mutant lines were stratified for 2 days in 4°C and sown in potting soil. The plants were grown under long day condition (16h light/8h dark) in a growth room at 21°C (light) or 19°C (dark). For water-loss assays on detached leaves, the rosette leaves of four-week-old WT and mutant plants were collected and placed in petri dishes. The dishes were left on a laboratory bench and weighted periodically. The assays were repeated at least three times.

For microscopic observation, the rosette leaves of four-week-old WT and mutant plants were collected and place in stomata opening buffer (20 mM KCl, 1 mM CaCl*_2_*, and 5 mM MES-KOH, pH 6.15) for 2 hours under light to open the stomata. We then used a tweezer to tear off the epidermal strips of the leaves and examined the stomata using a digital microscope (Hirox KH-7700). For ABA treatment, after being immersed in stomata opening buffer for 2 hours, the leaves were placed in a solution containing 1 μm ABA, and 2 hours later epidermal strips of the leaves were obtained and used for microscopic observation. The statistics of stomata, including stomata area, guard cell width at the center point (*W_middle_*), guard cell width at the quarter point (*W_quarter_*), stomatal width, and stomatal aperture were measured using ImageJ, and at least 6 visions were measured for each line. Sum of GCs’ width within a stoma was calculated as stomatal width minus stomatal aperture.

### Tissue-specific expression patterns of *MYS1* and *MYS2*

The promoter sequences of *MYS1* (2621 bp upstream of the translation starting codon ATG) and *MYS2* (2027 bp upstream of the translation staring codon ATG) were cloned into the TQ662 vector to drive the expression of the H2B-sGFP fusion reporter protein (Zhang *et al*., 2021). The vectors containing the *MYS1_pro_::H2B-sGFP* or the *MYS2_pro_::H2B-sGFP* fragment were transformed into *Agrobacterium tumefaciens* GV3101, which were then used to transform Arabidopsis WT plants by floral dipping (Clough and Bent, 1998). The GFP fluorescence of the positive transformed plants was detected with a confocal microscope (Zeiss LSM710).

### Subcellular localization of MYS1 and MYS2

For subcellular localization, the full-length genome sequences of GMS1 and GMS2 were fused upstream of GFP under the control of the super promoter in the pFK242 vector. The vectors were transformed into *Agrobacterium tumefaciens* ASE and then used to transform *Nicotiana benthamiana* epidermal cells via infiltration with a 1ml syringe. Plant leaves were cultured for 2 days and subjected to microscopic observation using a laser scanning microscope (Zeiss LSM880).

## Acknowledgments

We thank USTC Supercomputing Center and USTC School of Life Sciences Bioinformatics Center for providing the computing resources. This work was supported by grants from the Strategic Priority Research Program of the Chinese Academy of Science (XDA24010303), the National Natural Science Foundation of China (31770268), the Fundamental Research Funds for the Central Universities (WK2070000091), and University of Science and Technology of China (Start-up fund to S.M.).

## Author contributions

SM designed and supervised the project. YP, YL, FW, YQ, SM performed the experiment and analysis. YP, YL, and SM wrote the manuscript. All authors reviewed and approved the final manuscript.

## Supplemental Data

**Figure S1. UMAP plots of all cells from the three scRNA-seq datasets. (A, C, E)** UMAP plots of SHOOT1, ATML, and COTYLEDON. Colors indicate cell cluster ids. **(B, D, F)** Non-epidermis cells identified from the three datasets.

**Figure S2. Subcellular localization of MYS1 and MYS2 proteins.** MYS1-GFP and MYS2-GFP fusion protein driven by the 35S promoter were transiently expressed in *Nicotiana benthamiana* leaves and used for confocal microscopy observation. Bar, 50 μm.

**Figure S3. The sgRNAs and vectors used for generating *MYS1* and *MYS2* CRISPR knock-out mutant lines. (A)** The sgRNAs used for CRISPR knock-out vector construction. **(B)** The two vectors, pHY01 and pHY07, used for generating the CRISPR knock-outs. Both pHY01 and pHY07, as well as the method for cloning 1 or 2 sgRNAs into these two vectors, were developed in our lab previously (Geng *et al*., 2023), and they are shown here for illustration purpose. The *mys1-1* and *mys2-1* single mutants were generated using the pHY07 vector containing 1 sgRNA (sgRNA-300 or sgRNA-260). The *mys1 mys2-1* and *mys1 mys2-2* double mutants were generated using the pHY01 vector containing 2 sgRNAs (sgRNA-300 and sgRNA-260). The *mys1 mys2-3* and *mys1 mys2-4* mutants were generated using the pHY07 vector containing 2 sgRNAs (sgRNA-300-2 and sgRNA-260-2). Only T-DNA fragments are shown.

**Figure S4. *MYS1* and *MYS2* single mutants and their phenotypes. (A)** Sequencing results of *mys1-1* and *mys2-1* single mutants. **(B)** Water loss rate of detached leaves from *mys1-1*, *mys2-1*, and WT plants.

**Figure S5. Sequencing of *mys1 mys2-1* and *mys1 mys2-2* mutant lines.** Sequencing results of *mys1 mys2-1* (A) and *mys1 mys2-2* (B) mutant lines. Two sgRNAs, sgRNA-300 and sgRNA-260 contained within the same CRISPR construct, were used to target *MYS1* and *MYS2* respectively. “∼∼∼∼∼” indicates intron sequences.

**Figure S6. *mys1 mys2-3* and *mys1 mys2-4* double mutant lines and their phenotypes. (A, B)** Sequencing result of *mys1 mys2-3* and *mys1 mys2-4* mutant lines. Two sgRNAs, sgRNA-300-2 and sgRNA-260-2 contained within the same CRISPR construct, were used to target *MYS1* and *MYS2* respectively. “∼∼∼∼∼” indicates intron sequences. **(C)** Water loss rate of detached levees from *mys1 mys2-3*, *mys1 mys2-4*, and WT plants.

**Table S1. Gene co-expression modules related to stomata and epidermis extracted from AtGGM2014.**

**Table S2. Stage-specific gene expression and differential gene expression analysis during stomatal development and maturation.**

**Table S3. Sc_586: a list of genes up-regulated during stomatal maturation and function formation.**

**Table S4. Venn diagram analysis results of comparing sc_586 to other mature stomata-related gene lists.**

