## Supplemental Figures for "Stomatal maturomics: identifying genes regulating guard cell maturation and function formation from single-cell transcriptomes"

#### Fig. S1

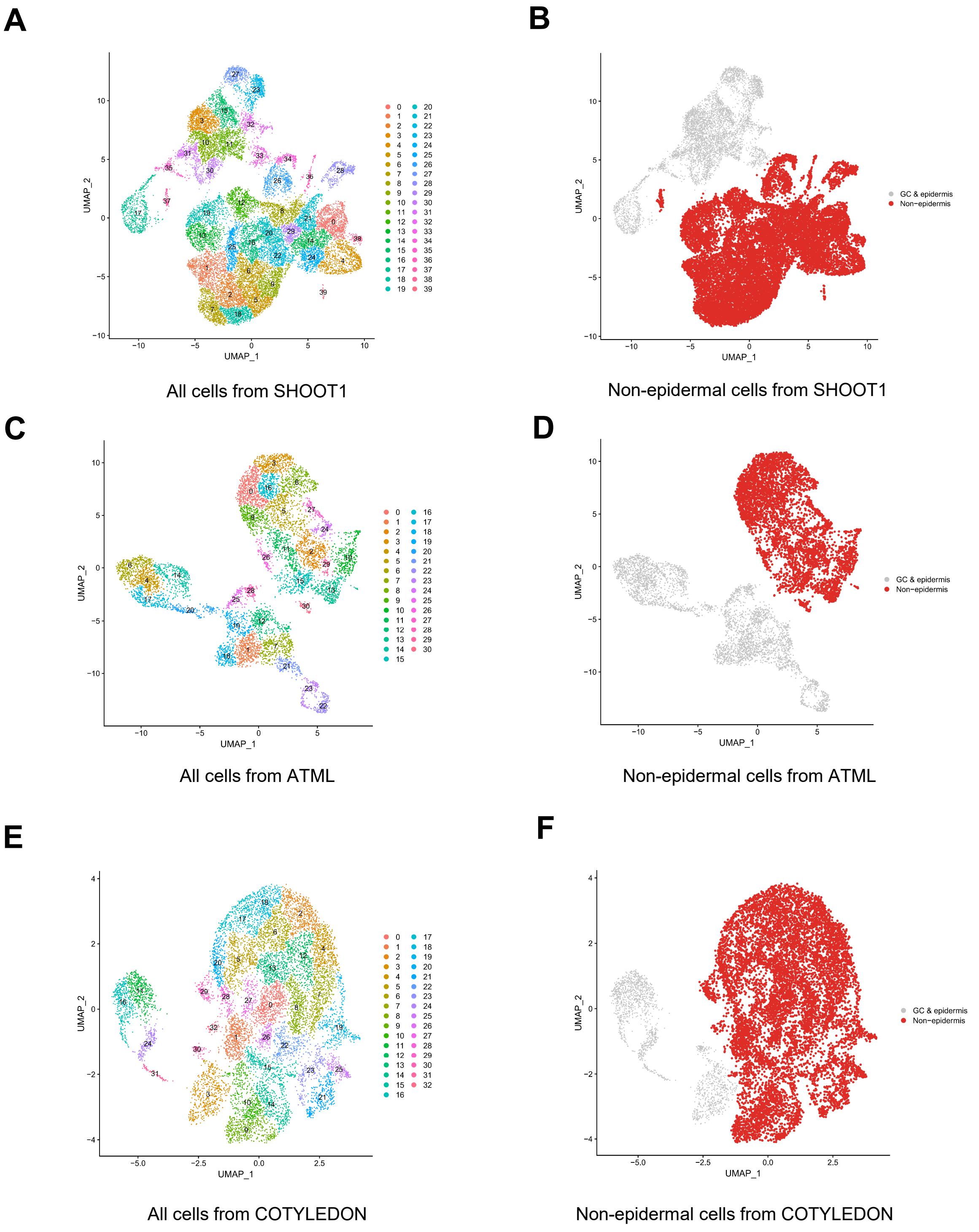


All cells from SHOOT1

Non-epidermal cells from SHOOT1

All cells from ATML

Non-epidermal cells from ATML

All cells from COTYLEDON

Non-epidermal cells from COTYLEDON

**Fig. S2**

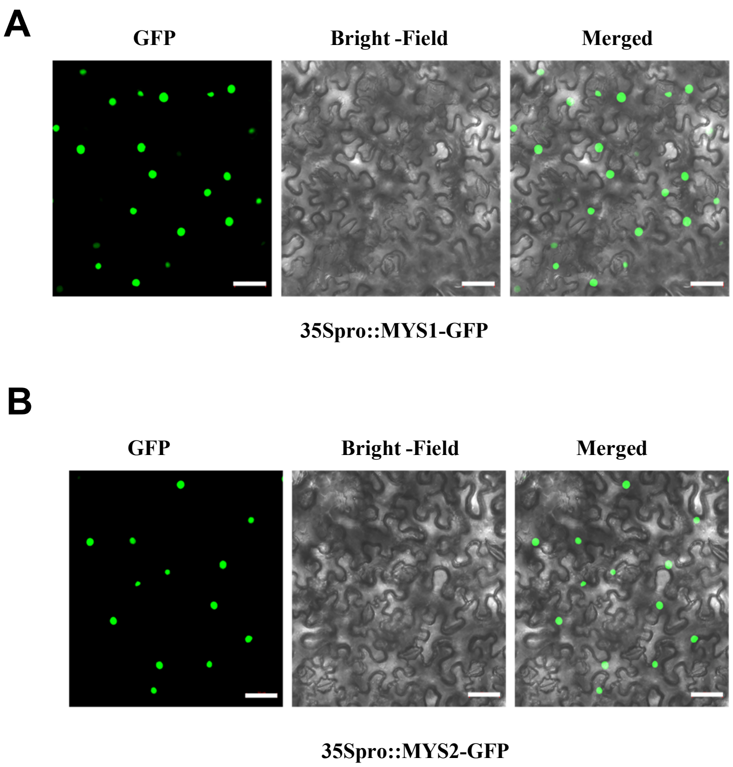
A
GFP
Bright -Field
Merged
35Spro::MYS1-GFP
B
GFP
Bright -Field
Merged
35Spro::MYS2-GFP

### Fig. S3

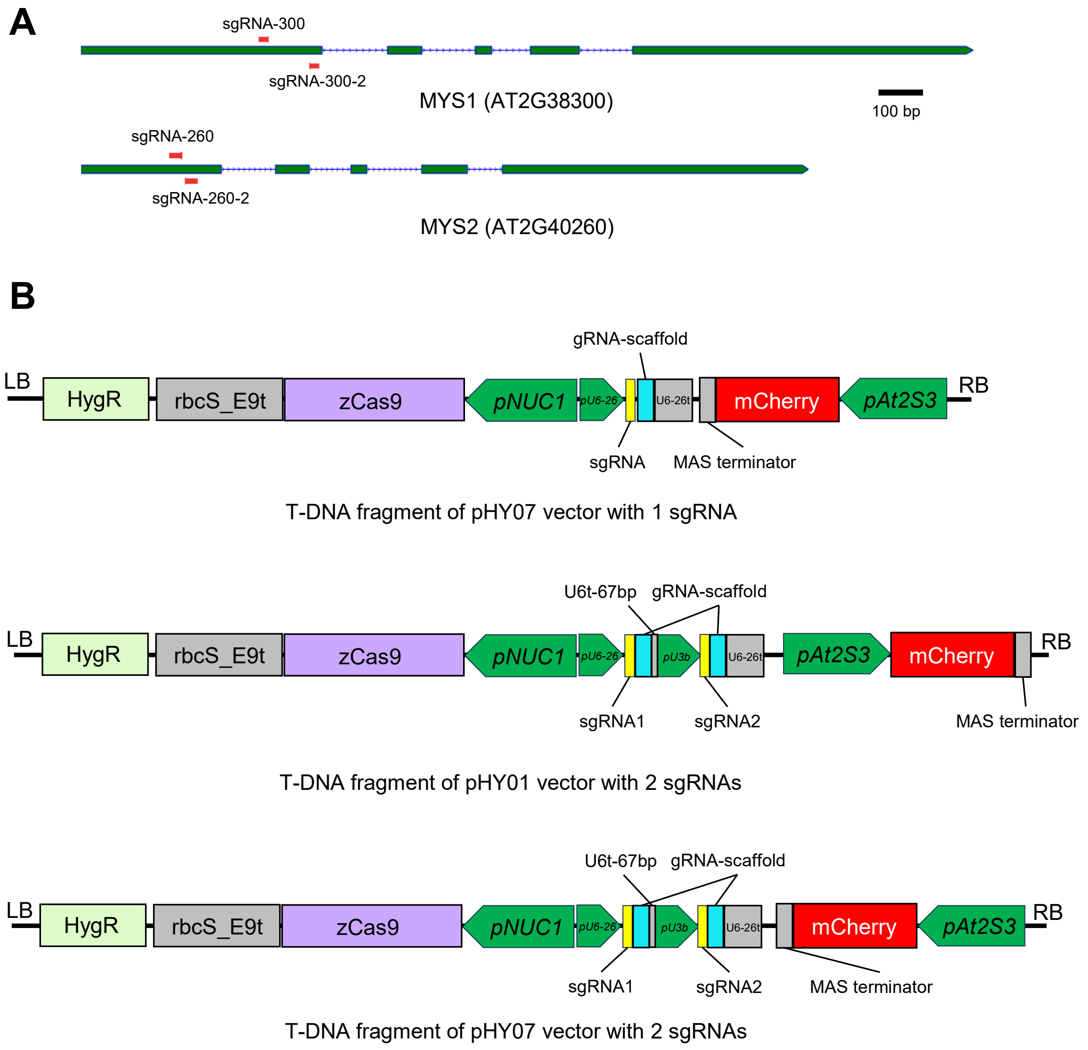
A
sgRNA-300
sgRNA-300-2
MYS1 (AT2G38300)
100 bp
sgRNA-260
sgRNA-260-2
MYS2 (AT2G40260)
B
gRNA-scaffold
LB
HygR
rbcS_E9t
zCas9
pNUC1
pU6-26
U6-26t
mCherry
pAt2S3
RB
sgRNA
MAS terminator
T-DNA fragment of pHY07 vector with 1 sgRNA
U6t-67bp
gRNA-scaffold
LB
HygR
rbcS_E9t
zCas9
pNUC1
pU6-26
pU3b
U6-26t
pAt2S3
mCherry
RB
sgRNA1
sgRNA2
MAS terminator
T-DNA fragment of pHY01 vector with 2 sgRNAs
U6t-67bp
gRNA-scaffold
LB
HygR
rbcS_E9t
zCas9
pNUC1
pU6-26
pU3b
U6-26t
mCherry
pAt2S3
RB
sgRNA1
sgRNA2
MAS terminator
T-DNA fragment of pHY07 vector with 2 sgRNAs

Fig. S4

A

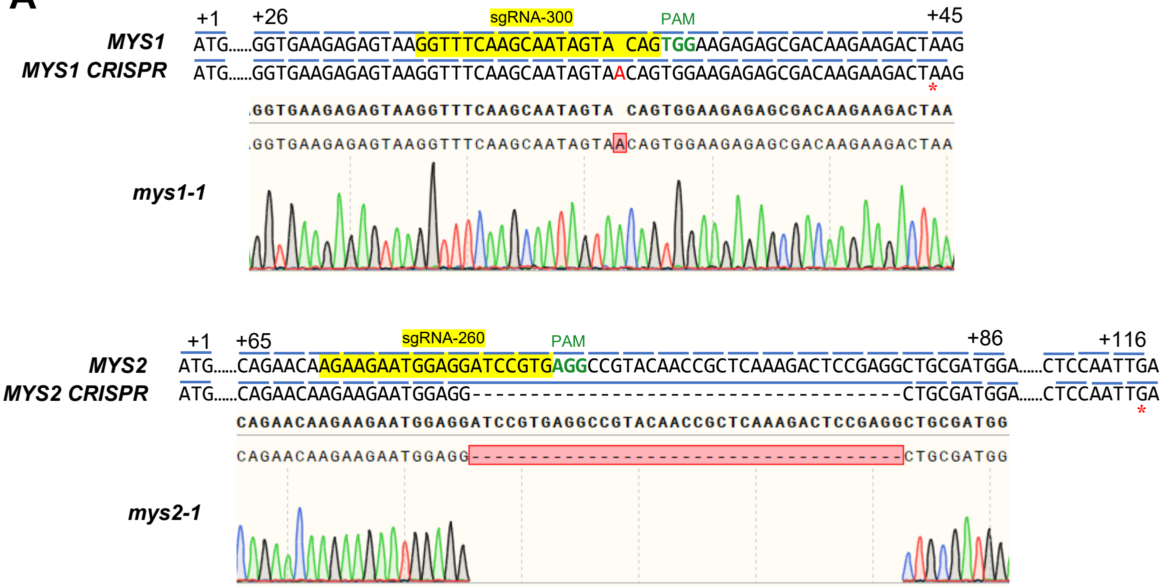
sgRNA-300
PAM
+1
+26
+45
MYS1
ATG......GGTGAAGAGAGTAAGGTTTCAAGCAATAGTA CAGTGGAAGAGAGCGACAAGAAGACTAAG
MYS1 CRISPR
ATG......GGTGAAGAGAGTAAGGTTTCAAGCAATAGTAACAGTGGAAGAGAGCGACAAGAAGACTAAG
mys1-1
sgRNA-260
PAM
+1
+65
+86
+116
MYS2
ATG......CAGAACAAGAAGAATGGAGGATCCGTGAGGCCGTACAACCGCTCAAAGACTCCGAGGCTGCGATGGA......CTCCAATTGA
MYS2 CRISPR
ATG......CAGAACAAGAAGAATGGAGG-----------------------------------CTGCGATGGA......CTCCAATTGA
mys2-1

B

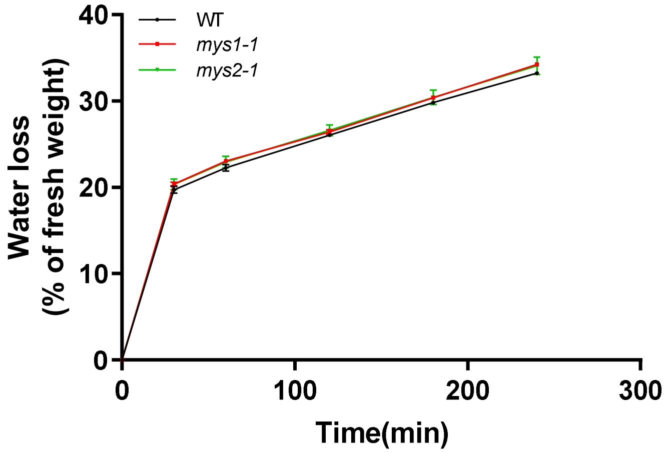
WT
mys1-1
mys2-1
Water loss
(% of fresh weight)
Time(min)
0
10
20
30
40
0
100
200
300

### Fig. S5

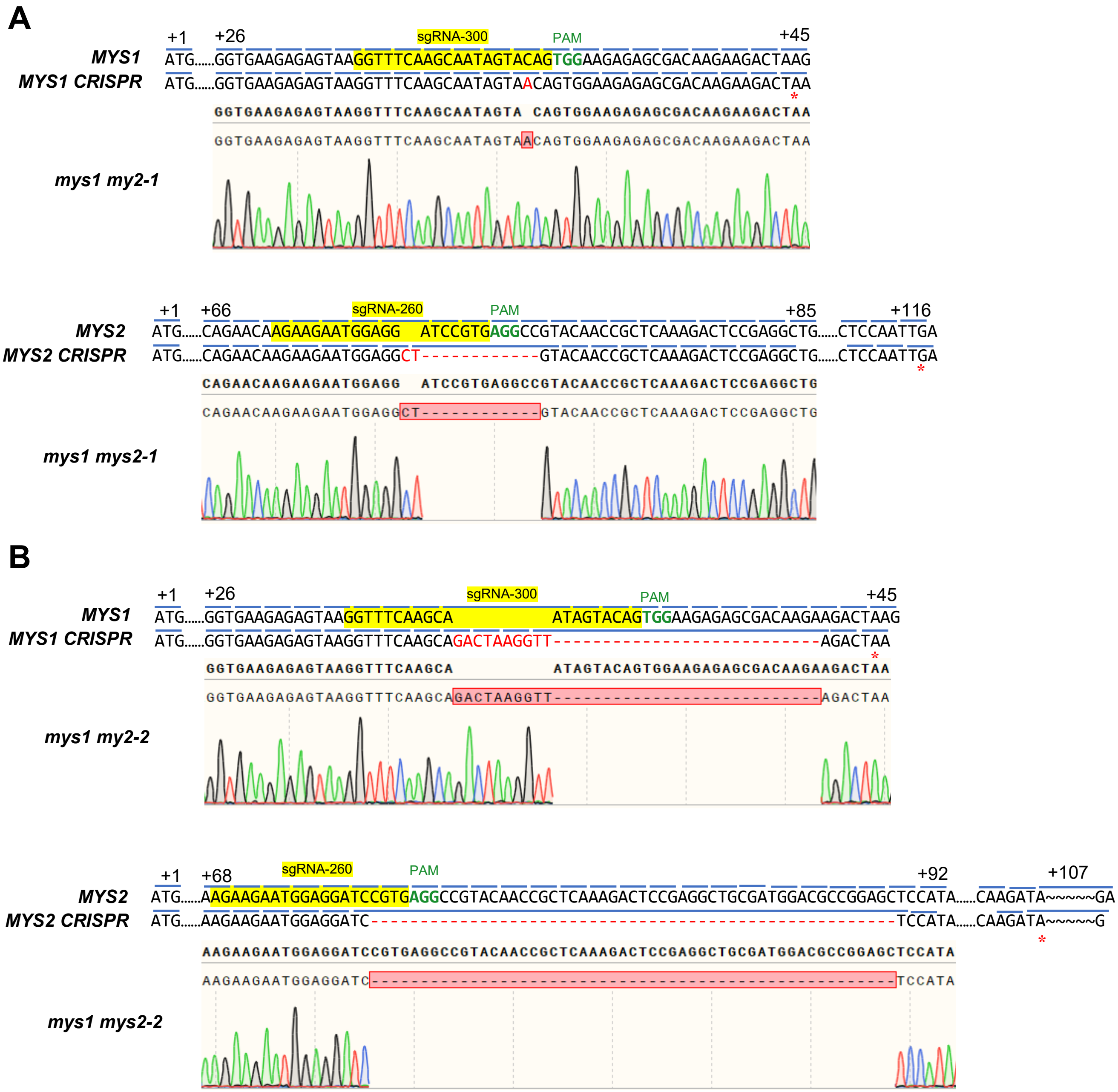
A
+1
+26
sgRNA-300
PAM
+45
MYS1
ATG......GGTGAAGAGAGTAAGGTTTCAAGCAATAGTACAGTGGAAGAGAGCGACAAGAAGACTAAG
MYS1 CRISPR
ATG......GGTGAAGAGAGTAAGGTTTCAAGCAATAGTAACAGTGGAAGAGAGCGACAAGAAGACTAA
GGTGAAGAGAGTAAGGTTTCAAGCAATAGTA CAGTGGAAGAGAGCGACAAGAAGACTAA
GGTGAAGAGAGTAAGGTTTCAAGCAATAGTAACAGTGGAAGAGAGCGACAAGAAGACTAA
mys1 my2-1
+1
+66
sgRNA-260
PAM
+85
+116
MYS2
ATG......CAGAACAAGAAGAATGGAGG ATCCGTGAGGCCGTACAACCGCTCAAAGACTCCGAGGCTG......CTCCAATTGA
MYS2 CRISPR
ATG......CAGAACAAGAAGAATGGAGGCT------------GTACAACCGCTCAAAGACTCCGAGGCTG......CTCCAATTGA
CAGAACAAGAAGAATGGAGG ATCCGTGAGGCCGTACAACCGCTCAAAGACTCCGAGGCTG
CAGAACAAGAAGAATGGAGGCT------------GTACAACCGCTCAAAGACTCCGAGGCTG
mys1 mys2-1
B
+1
+26
sgRNA-300
PAM
+45
MYS1
ATG......GGTGAAGAGAGTAAGGTTTCAAGCA ATAGTACAGTGGAAGAGAGCGACAAGAAGACTAAG
MYS1 CRISPR
ATG......GGTGAAGAGAGTAAGGTTTCAAGCAGACTAAGGTT--------------------------AGACTAA
GGTGAAGAGAGTAAGGTTTCAAGCA ATAGTACAGTGGAAGAGAGCGACAAGAAGACTAA
GGTGAAGAGAGTAAGGTTTCAAGCAGACTAAGGTT--------------------------AGACTAA
mys1 my2-2
+1
+68
sgRNA-260
PAM
+92
+107
MYS2
ATG......AAGAAGAATGGAGGATCCGTGAGGCCGTACAACCGCTCAAAGACTCCGAGGCTGCGATGGACGCCGGAGCTCCATA......CAAGATA~~~~~GA
MYS2 CRISPR
ATG......AAGAAGAATGGAGGATC-----------------------------------------------------TCCATA......CAAGATA~~~~~G
AAGAAGAATGGAGGATCCGTGAGGCCGTACAACCGCTCAAAGACTCCGAGGCTGCGATGGACGCCGGAGCTCCATA
AAGAAGAATGGAGGATC-----------------------------------------------------TCCATA
mys1 mys2-2

#### Fig. S6

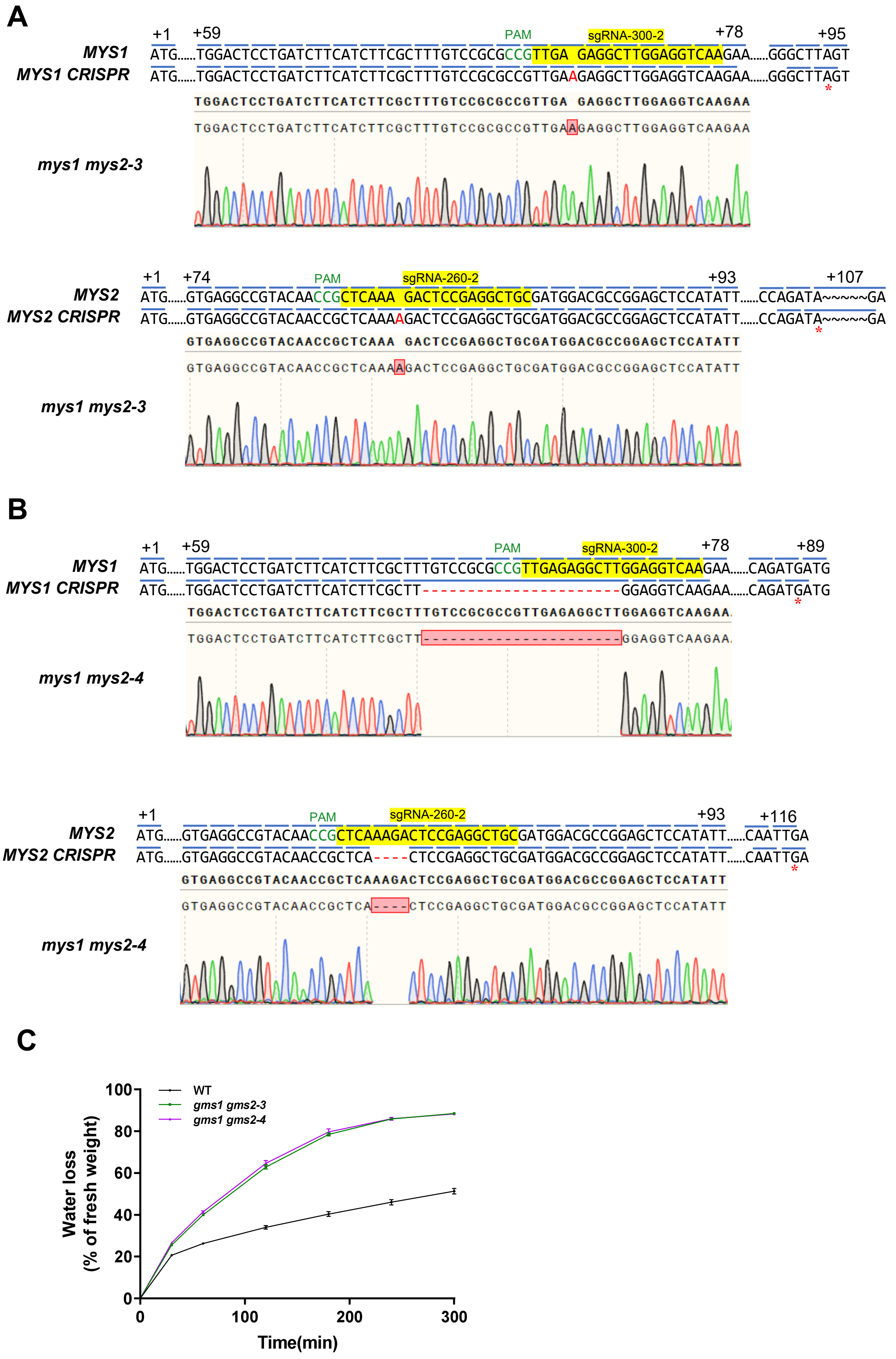

A
MYS1
MYS1 CRISPR
PAM
sgRNA-300-2
+1
+59
+78
+95
mys1 mys2-3
MYS2
MYS2 CRISPR
sgRNA-260-2
+74
+93
+107
B
+89
+116
mys1 mys2-4
C
WT
gms1 gms2-3
gms1 gms2-4
Water loss
(% of fresh weight)
Time(min)
0
20
40
60
80
100
100
200
300
